# Intrapopulation adaptive variance supports selective breeding in a reef-building coral

**DOI:** 10.1101/2021.05.21.445206

**Authors:** Crawford Drury, Nina Bean, Casey Harris, Josh Hancock, Joel Hucekba, H. Christian Martin, Ty Roach, Robert Quinn, Ruth D. Gates

**Affiliations:** Hawaiʻi Institute of Marine Biology, University of Hawaiʻi, Kāneʻohe, HI, USA; University of Amsterdam, Amsterdam, Netherlands; Department of Biochemistry and Molecular Biology, Michigan State University, East Lansing, MI, USA

**Keywords:** Coral Bleaching, Genetic Association Analysis, Metabolomics

## Abstract

The long-term persistence of coral reefs under climate change requires heritable drivers of thermal tolerance which support adaptation. The genomic basis of thermal tolerance has been evaluated across strong spatial and environmental gradients, but this variation also exists within populations due to neutral evolutionary processes. Small scale heterogeneity in coral bleaching is ubiquitous, so we used corals from a single reef to examine genomic signatures of bleaching performance, their biochemical correlates and the downstream consequences of selective breeding. In the absence of directional selection due to environmental differences, adult corals from a single population exhibit strong genomic patterns related to natural bleaching tolerance and symbiosis state, including functional differentiation in signaling pathways, protein and amino acid modification and metabolism. Conversely, growth, development and innate immune responses did not distinguish bleaching tolerance in adult corals. The genomic signatures of these gene ontologies influence biochemical patterns in healthy corals, primarily via cell-signaling pathway impacts on peptides and amino acids. Thermal tolerance in this population is highly heritable, with significantly higher survivorship under temperature stress in larvae and juveniles reared from thermally tolerant parents than those from sensitive parents. Using a select and re-sequence approach, certain gene ontologies were reproducibly impacted, while antioxidant activity and cell signaling ontologies were disproportionately selected in thermally tolerant corals, demonstrating the genomic drivers of successful selective breeding. These data show that intrapopulation variance in the absence of historical selection supports the adaptive capacity of coral reefs under climate change.

## Introduction

Reef-building corals are complex holobionts composed of a cnidarian host, symbiotic dinoflagellates in the family *Symbiodiniacea* and a suite of other micro-organisms. Each compartment of the holobiont interacts to dictate trait values like thermal tolerance, which is increasingly important for ecosystems facing climate change. Corals are susceptible to heat stress, which leads to the breakdown of the coral-algal symbiosis and compromises the structure and function of reefs ^1,2^. Despite these threats, there is substantial variation in thermal tolerance within and among coral populations that can exist over very small spatial scales ^3–9^ and supports the adaptive potential of reef-building corals ^10–12^.

Genomic variation is an important driver of coral bleaching ^3,13,14^, but previous research on the adaptive capacity of the host has focused almost exclusively on differences driven by strong selection across obvious environmental gradients ^5,13–17^. This framework has also been leveraged in breeding experiments ^12,14^, limiting our understanding of the adaptive potential of reefs over scales such as single reefs or bays, where fertilization is most likely to occur. Nested within more subtle environmental contours, variation in coral bleaching produced by other demographic processes is ubiquitous and is the raw material upon which evolution acts within a coral metapopulation. This mismatch represents a mostly unknown reservoir of resilience in the absence of *a priori* environmental correlates that may be increasingly important as reefs become more isolated in climate refugia^18^.

*Montipora capitata* is a cosmopolitan reef-building coral that is dominant in Kāneʻohe Bay, Oʻahu, Hawaiʻi. *M. capitata* is a vertically transmitting broadcast spawner which releases symbiont provisioned eggs, creating a tight link between host and symbiont ^19,20^ which may facilitate ‘adaptive’ change by diversifying sources of resilience. Bleaching phenotype in this *M. capitata* population is related to symbiosis with *Cladocopium* or *Durusdinium* ^21,22^, where individual corals typically harbor *Cladocopium* and are more thermally sensitive (*‘*bleached’ phenotype) or a mixed community of both genera and are more thermally tolerant (‘nonbleached’ phenotype). There are infrequent exceptions to these patterns^21^, but these relationships appear broadly fixed through time and multiple bleaching events^22^, suggesting that a tight connection between host genotype and symbiont constituency does not allow for symbiont ‘shuffling’. Nested within these patterns, there is a strong genotype effect on the biochemical signature of metabolites in these corals, including those generated by the symbionts ^23^, suggesting that the coral host either actively or passively affects symbiont biochemistry *in hospite*.

We use this framework to examine intrapopulation genomic variance and the consequent biochemical signatures that define coral thermal tolerance. We then bulk cross gametes from each parental coral phenotype, documenting improved thermal tolerance in larvae and juveniles from tolerant parents, and use a select and re-sequence approach to define genomic variants associated with improved thermal tolerance in larvae.

## Methods

### Collections

In 2015, the summer thermal maximum produced a mosaic of bleaching in Kāneʻohe Bay, where adjacent colonies either severely bleached or remained visually healthy ^21,24,25^. The majority of these colonies recovered and have been tracked through time (Fig. S1). We selected 22 healthy colonies (11 pairs) along a 2-3m depth contour on Reef 13 in Kāneʻohe Bay and used them for genetic, metabolomic and gamete collections for selective breeding. We collected 3 replicate branches for adult genetics and symbiont extractions in August 2018 (to avoid spawning impacts) and collected gamete bundles from each colony *in situ* on 13 July 2018. We collected metabolite samples from each colony in May-September 2019^23^.

### Crosses, Fertilization and Treatments

All 22 colonies in the study spawned on 13 July 2018. We created a site-wide cross (gametes from all-colonies in the study), bleached phenotype pool (only gametes from historically bleached parents) and nonbleached phenotype pool (only gametes from historically nonbleached parents) prior to the breakup of bundles and fertilization (Fig. S1). Gamete bundles from each bulk cross were immediately aliquoted (n=10) for fertilization^26^ and returned to the Hawaii Institute of Marine Biology. Triplicates of fertilized embryos from each phenotype were added to one 150L larval rearing conical and allowed to develop for 12 hours to the prawn-chip stage before temperature treatments (ambient: 27.4 ± 0.55°C; mean ± 1 SD) and high: 30.2 ± 0.51°C) began for each of three phenotype pools (Fig. S2).

### Larval Survivorship, Juvenile Settlement and Survivorship

We established replicate survivorship trials at 12 hours post fertilization (hpf; n=15 per phenotype and temperature) and measured larval survivorship at 27, 47, 75, 96, 120, 172 and 213 (hpf) (Fig. S2). At 109hpf, larvae from each phenotype at ambient temperatures were transferred to settlement chambers (n=4) and allowed to settle on pre-conditioned aragonite plugs. After 8 days exposure to substrate, plugs (n=877) were randomly allocated into ambient (27.6 ± 0.96°C) and high (30.1 ± 0.38°C) temperature treatments (n=2 per temperature) (Fig. S3).

Settlement plugs were photographed at the initiation temperature treatments (12 days post fertilization) and at 8, 14, 18, 27 and 34 days in treatments. Individual settlers were counted from photos at each timepoint. The removal of larvae for settlement did not interrupt the ongoing larval survivorship trials. See Fig. S2 for details.

Larval and juvenile survivorship was analyzed with Kaplan-Meier regression in the R package *Survival*, with pairwise comparisons between temperature and phenotype crosses using a likelihood ratio test (implemented by *Survminer*::*pairwise_survdiff*) with Bonferroni adjusted p-values. We calculated heritability of both larvae and juveniles separately at each temperature, excluding the cross phenotype. We defined broad-sense heritability as variance (MSE) explained by Phenotype in a 1-way ANOVA as a proportion of total variance (MSE), modified from Falconer ^27^.

### Symbiont Analysis

Larval symbiont collections were made at 16, 36, 60 and 84 hours post fertilization, representing daily sampling that overlapped genomics sampling (Fig. S2). Three replicates of 30-50 larvae were haphazardly collected from each treatment. Adult fragments and larval timeseries collections were extracted with a CTAB-chloroform protocol (following^22^). Concentration of *Cladocopium, Durusdinium* and the coral host were quantified via quantitative PCR using actin and Pax-C assays ^21,28,29^ on a StepOnePlus system (Applied Biosystems) with two technical replicates. Samples were re-extracted and analyzed if C_T_ replicate standard deviation was greater than 1 between replicates or if only a single replicate amplified. C_T_ values were corrected for fluorescence and copy number using the equations of ^21^. See Fig. S2 for details.

### Library Prep and Sequencing

Adult fragments were stored at −20°C until processing. Larval samples were collected from each conical (n=6) at 16 hpf by collecting 5 replicates of 50 larvae (n=250 total larvae per timepoint per treatment), preserved in 2% SDS in DNAB and stored at −20°C until processing. A second sample was collected in the same way at 84 hpf, representing timepoints near the initiation of heat stress during late development and after the putative selective pressure had led to ∼25% mortality (Fig. S2).

DNA was extracted using a DNeasy Blood & Tissue Kit (Qiagen) following manufacturer protocol. Extracted DNA was quantified with a Qubit fluorometer and 300ng of DNA was digested with *HindIII* (New England Biolabs). Libraries (n=23; 22 adult samples and an adult technical replicate, 60 larval samples) were prepared with an Illumina TruSeq Nano kit and TruSeq adapters for 350bp inserts following manufacturer protocol. Samples were sequenced on a single lane of an Illumina HiSeq 4000 with paired end 150bp reads (Genewiz, South Plainfield, NJ).

We prepared a reference genome by mapping the *Montipora capitata* genome to the chromosome scale scaffolding of *Acropora millepora ^13^* using default options in RagTag ^30^, which resulted in ∼5% higher alignment rates and higher contiguity. The *Montipora capitata* individual used for genomic sequencing showed signals of gene duplication ^31^ and is ∼885MB (compared to 458 MB for the *A. millepora* reference). We included un-anchored *M. capitata* sequences as additional contigs, effectively increasing the contiguity of the reference without removing any data. Short read alignment rates were substantially lower against the original *A. millepora* genome or against the chromosome scale *M. capitata* anchored alignment without additional contigs. We annotated predicted protein sequences from the *M. capitata* assembly ^31^ using blastp against Uniprot/Swissprot and extracted gene ontologies using a custom script. We used annotations from the *A. millepora* assembly ^13^ for scaffolded contigs and the additional annotations for unscaffolded *M. capitata* sequences using a custom R script.

### Adult Analysis

Raw reads were demultiplexed at the sequencing facility and we trimmed adapters using *cutadapt*^32^ before aligning all samples using *bwa mem*^33^. We analyzed adult clonality using a biological replicate in ANGSD^34^ to generate an identity by state matrix with filtered reads (minMapQ 30, minQ 30) and used hierarchical clustering to confirm there were no clonemates in our adult population. We then examined the association of individual loci with each adult phenotype (n=21, one sample removed with low depth) using genotype probabilities in ANGSD (GL 2, doAsso 1, doGeno 8, doPost 1, minDepth 5, SNP pvalue<2e^−6^). We permuted phenotype assignments 10,000 times, recalculated association and selected the highest LRT value for each permutation, then assigned the 99^th-^ percentile of this distribution as the cutoff for significance (p<0.01) for individual loci. We used *PCAngsd*^35^ and *NGSadmix*^36^ to examine population structure with default settings.

We used annotations within 2.5kbp of each variant to derive ontologies for GO_MWU^37^, treating the LRT statistic from the association as ‘heats’ for all loci (n=20,409). This strategy resolves gene ontologies that are enriched in high or low association values, representing functional groupings that are different between phenotypes or are significantly similar and do not distinguish bleaching tolerance. We extracted all loci annotated with each significant ontology (FDR p<0.1) and ran a *PCAngsd* subroutine with these loci to summarize genetic differentiation in each functional category.

### Larval Analysis

To evaluate selection in the larval resequencing experiment, we created alignment files of pooled larval replicates by treatment, phenotype and timepoint and pooled adults by phenotype (see Table S1 for details). We summarized allele counts using *bcftools* and exported biallelic variants. We analyzed per-site F_ST_ for each sample (depth>=10) using the R package *poolfstat* (v1.1.0) ^38^ and calculated a Population Branch Statistic (PBS)^39^, which provides information on divergent selection between multiple populations (see Table S1 for details). There was low and uneven coverage in the bleached samples, so we excluded this pool from downstream analysis.

In this context, the PBS statistic is ideal for low-depth data because large values are derived from cases where secondary alleles are found at similar frequencies in both the parental pool and the initial larval pool, assuring that signals of heterozygosity are not a sequencing artifact. PBS of the comparison of interest is also positively influenced by similarity in the ‘outgroups’, in this case the initial larval and the parental pool. This expected pattern (based on random crosses from a pool of known parental genotypes), further supports consistent sequencing outcomes, minimizes artifacts and highlights selection. We further filtered the dataset to loci where both alleles were found in parental and initial pools and where the initial-parental F_ST_ was less than 0.05.

To ask how heat selection impacted specific loci, we calculated a PBS statistic distinguishing the final post-heat larval pool from the initial and parental larval pools for each phenotype. To evaluate selection patterns, we used GO_MWU ^37^, treating the PBS statistic as ‘heats’ for all loci shared between the nonbleached and cross populations (n=31,794 loci). We repeated this analysis on larvae from ambient tanks (n=26,407 loci) to confirm that heat-selected ontologies were not due to off-target mortality effects and documented 2 overlapping ontologies (ossification, circadian sleep/wake cycle process), which were removed.

We also calculated the difference between PBS in nonbleached and site-wide cross larvae and used the values for GO_MWU enrichment analysis. This framework isolates functional categories that are disproportionately selected in nonbleached corals and contribute to selective breeding outcomes.

### Metabolite Analysis

Small biopsies of all 22 parent corals were extracted in 500uL 70% methanol, kept on ice for 30 minutes and stored at −80°C until shipping to Michigan State University. Methanolic extracts were analyzed using LC-MS/MS. Details of chromatography and mass spectrometry parameters can be found in ^23^. The converted .mzXML files were processed with MZmine2^40^ and the feature quantification table was exported and processed with the Global Natural Products Social Molecular Networking (GNPS)^41,42^ and CANOPUS on SIRIUS 4^43,44^. MZmine2 parameters for processing mass spectrometry data are described in^23^, and the CANOPUS parameters can be found in Table S2. Both CANOPUS output files (“canopus_summary.csv”,”compound_identification.csv” were merged and filtered to include the most specific class of molecules, matches and their molecular formulas. Molecular features from blanks and found between both files that did not display *in silico* annotations from CANOPUS and correlation between molecular formulas were removed.

We repurposed the WGCNA analytical framework for metabolites, which summarizes molecules with highly correlated abundances into modules^45^. We used log2 transformed abundances after imputing missing data with the lowest abundance of each molecule following ^46,47^. We tested a power threshold from 1 to 20 and chose 6 as the ‘elbow’ in scale-free topology, with modules merged > 0.15, minKME = 0.8, at least 25 molecules per module and an unsigned network. We used the first principal component from *PCAngsd* as a summary of the genetic position of each coral’s genes relating to that ontology as the ‘trait’ for Spearman correlation analysis with WGCNA. We used a Fisher’s exact test to test for enrichment in molecular families within each module and calculated a false discovery rate-corrected p-value.

### Betaine Analysis

After noting the high association and strong linkage between two loci of interest, we extracted a 50kb window centered at the midpoint of these loci and annotated with blastn against cnidarians. We documented glycine sarcosine dimethylglycine N-methyltransferase within this selection window, a gene family that produces Betaine Glycine (BG), which we annotated in the metabolite data (metabolomics standards initiative level two^48^). We compared Betaine relative abundances between phenotypes using a Wilcox test. We used the raw annotations from GO_MWU (before merging into the parent term ‘Cellular Modified Amino Acid Metabolic Process’), and compared the association values for ‘Amino-Acid Betaine Metabolic Process’ to all other loci using a Wilcox test.

## Results

We used a global analysis of genetic data to examine clonality and the underlying proportion of historical bleaching phenotype determined by the host. Corals in this study were unique genotypes, but bleaching phenotype (and by extension symbiont constituency) was not resolved by genotype hierarchical clustering (Fig. 1a). Principal component analysis of genotype likelihoods shows strong genomic patterns underlie bleaching phenotype (PERMANOVA p=0.003, R^2^ =0.24, Fig. 1b) within a single population (*NGSadmix* K=1), explaining 24% of variance. Interestingly, two individual corals clustered with the opposite phenotype (Fig. 1b). We also evaluated symbiont community in the parents to provide additional context for phenotype. Corals with the nonbleached phenotype were dominated by *Durusdinium*, frequently with background levels of *Cladocopium* at <10% abundance. Conversely, all Bleached phenotype corals harbored *Cladocopium*, except colony 11 which contained *Durusdinium* in 2 replicates. Replicate sampling within colonies suggests occasional heterogeneity in symbiont type (Fig. 1c).

**Figure 1.**
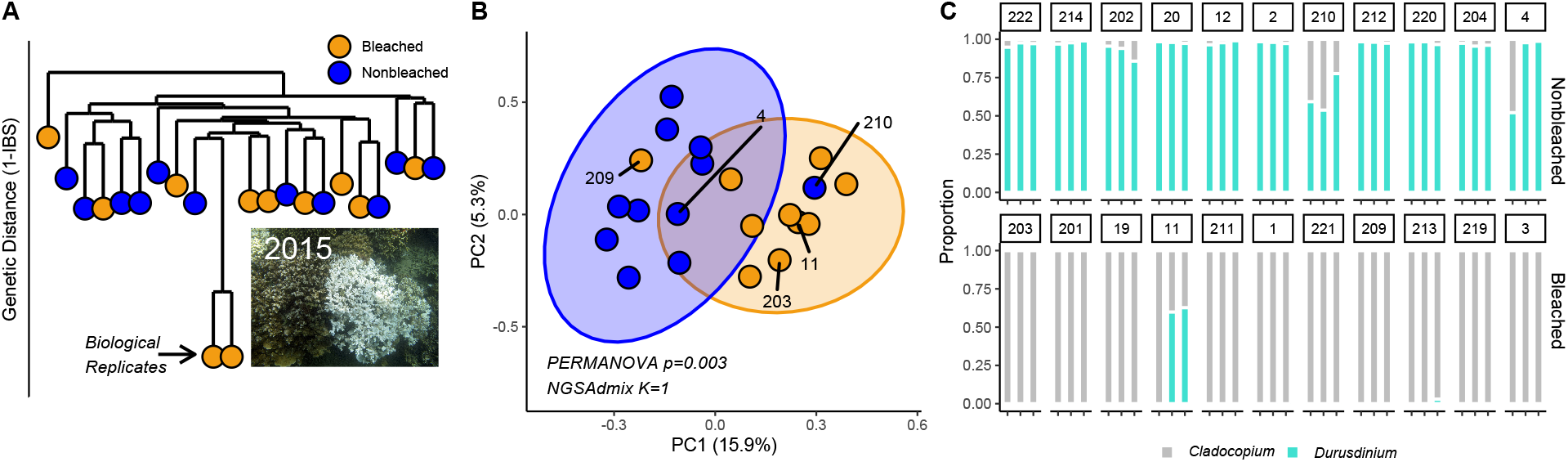
Genomic and symbiotic dynamics underly bleaching phenotype in Montipora capitata. A) Hierarchical clustering of genetic distances calculated from genotype likelihoods, with one sequenced pair of biological replicates to confirm clonality. Inset is example historical bleaching phenotype from 2015. Corals were fully recovered at the time of this experiment. B) PCA of 20,407 loci, shaded by historical bleaching phenotype. C) Symbiont communities in three haphazard replicates of each colony used in this study, shaded by proportion of *Cladocopium* or *Durusdinium* measured via qPCR.

We used association testing of genotype likelihoods to determine loci that were significant drivers of phenotype and used permutation testing (n=10,000) to generate p-values for the outcome of the association test. Thirty-six loci were significantly associated with phenotype (p<0.01, Table S3), but we detected no private alleles in these loci and no loci were fixed; all contrasts were between the heterozygote and homozygote, similar to ^14^. There were several individual loci near genes previously implicated in thermal tolerance, including plexin-B like, D-inositol 3-phosphate glycosyltransferase-like, kelch-like protein diablo, Mucin-like, monocarboxylate transporter 10-like, and several protein ubiquitin ligases. Due to the sparsity of SNP data, these loci are unlikely to be causative, so we used GO_MWU to analyze functional categories which are significantly enriched in loci that differentiate phenotypes.

We used the LRT values from adult association testing for 20,407 loci as ‘heats’ to find functional groups corresponding to phenotype. Several ontologies (n=28) were enriched in loci with high association values, suggesting a significantly elevated influence on phenotype compared to background. These ontologies included protein metabolism, localization and modification, GTPase activity, ribosomal structure, cell-signaling, exopeptidase activity, Erk cascades and endosomal membranes (Fig. 2a; Table S4). Conversely, steroid metabolism, innate immune response, developmental regulation, cell cycling, downregulation of cell growth, the endoplasmic reticulum, ds-DNA binding, coreceptor binding and calcium ion binding were enriched in low LRT values, suggesting a significant similarity between phenotypes in certain functions/processes that do not influence thermal tolerance (n=48 ontologies; Table S5).

**Figure 2.**
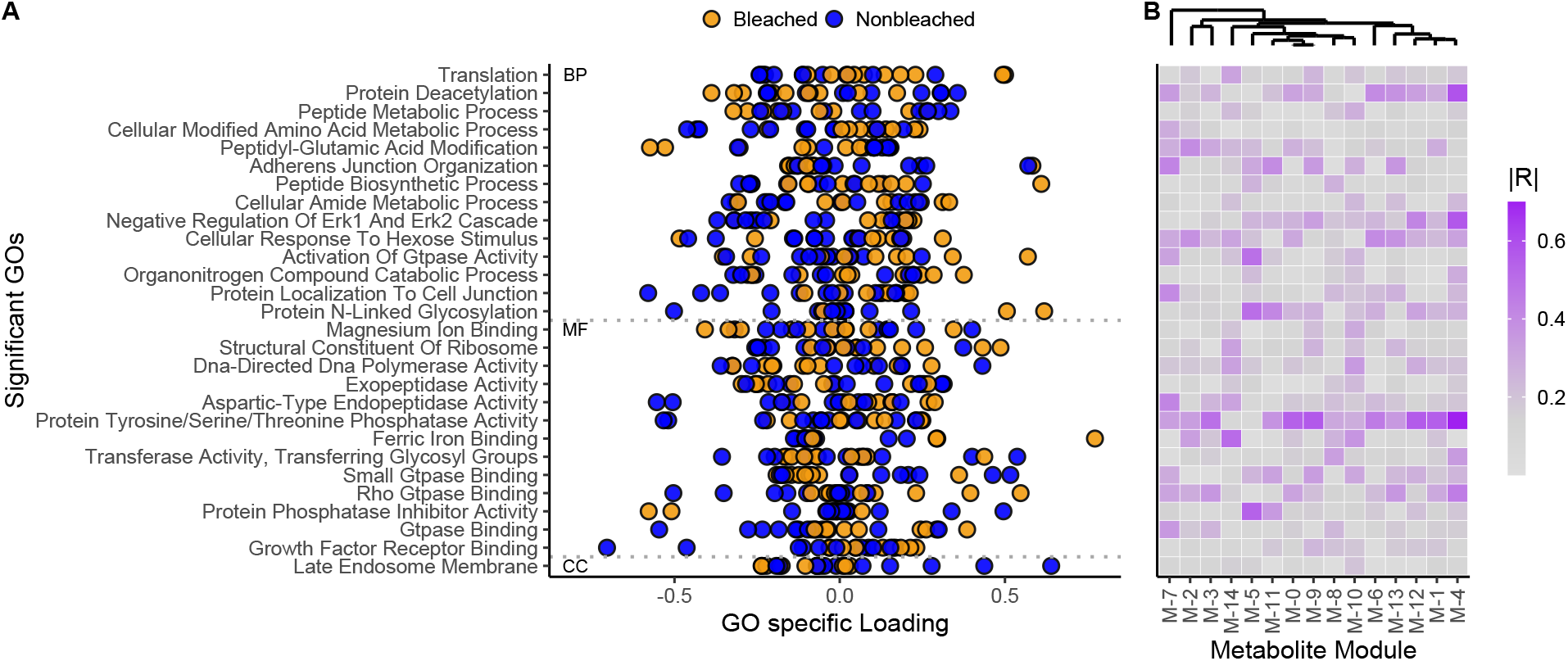
Functional ontologies distinguish bleaching phenotype and relate to biochemical patterns. A) The 28 gene ontologies that were significantly enriched in loci highly associated with bleaching phenotype. Points represent each colony in the study along the first principal component that describes all loci associated with the ontology of interest. B) WGCNA modules of the corals in this study, shaded by correlation with loading in (A). Each column represents a metabolite module resolved by WGCNA and each row corresponds to gene ontologies to the left. No values were significant after FDR correction.

Multilocus variation related to Protein Deacetylation, Erk Cascade Regulation and Protein Tyrosine/Serine/Threonine Phosphatase activity broadly impacted the biochemistry of corals in this study (Fig. 3). There were high correlations between metabolite Module-4 and each of these gene ontologies (R>0.5, p<0.0005). While these correlations were not significant after multiple comparisons correction, the conserved response of M-4 with multiple phenotype-defining gene ontologies suggests that this pattern is biologically important. Module-4 was significantly enriched in peptides, dipeptides, N-acyl-alpha amino acids and Trifluoromethylbenzenes (FDR p<0.03). Module-5 was also highly correlated (R>0.5) with Protein Phosphatase Inhibition Activity, Protein N-linked Glycosylation and GTPase activity and significantly enriched in menaquinones and prenol lipids (FDR p<0.02). The betaine lipid identified in ^23^, an important molecule for distinguishing phenotype in this species, was in Module-7.

**Figure 3.**
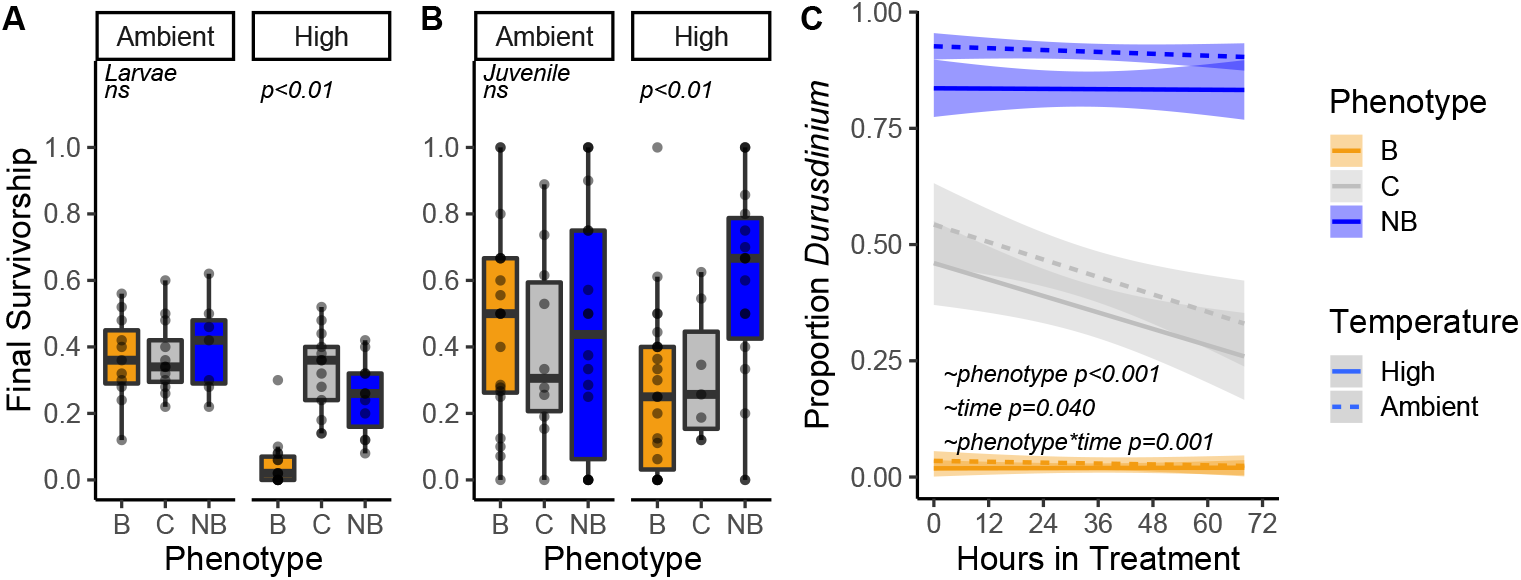
Selective Breeding Dynamics in Larval and Juvenile corals. A) Final survivorship 213 hours post fertilization for larvae at ambient and high temperatures. B) Final survivorship 34 days post settlement for juveniles at ambient and high temperatures. Boxplots are mean 1± IQR. C) Proportion *Durusdinium* of the three phenotype larval pools from fertilization to 84hpf in ambient and high temperatures. Line is best-fit with 95% confidence interval.

Among the top 36 individual loci for defining phenotype (LRT permutation p<0.01), two were on Chromosome 11 separated by approximately 11kb (Chr11_4203672 and Chr11_4215008). These loci were strongly linked (R^2^ =0.87), suggesting coinheritance and selection on nearby genes. We annotated a 50kb window centered on the variants of interest (following ^14^) using blastn. Several predicted genes were within this window, including PAX-c (e=3e^−104^), histone-lysine N-methyltransferase 2E-like (e=7e^−176^), leucine-rich repeats and immunoglobulin-like domains protein 1-like (e=5e-^115^), octopamine receptor beta-2R-like (e=1e-^109^), MAATS1-like protein (e=4e^−112^) and glycine sarcosine dimethlycine N-methylransferase-like (GSMT/SDMT, e=2e-^105^).

The product of GSMT/SDMT is Betaine Glycine (BG), which we documented in our metabolite dataset. BG was more abundant in nonbleached corals than their bleached counterparts (Wilcox test p=0.005; Fig. S4); however, colony 203 was an extreme outlier in the bleached phenotype. If this colony was excluded, concentrations were 22.8% higher in Nonbleached corals (Wilcox test p<0.001). We examined loci annotated as ‘Amino-acid Betaine Metabolic Process’ (GO: 0006577, n=10) with a Wilcox test and found a significantly higher LRT than other ontologies (p=0.013), suggesting this ontology is enriched in loci that discriminate between bleaching phenotypes.

To assess the adaptive potential of this intrapopulation genetic variation, we selectively crossed gametes from the parents on 13 July 2018, when all 22 colonies in the study spawned. A total of 11 colonies of each phenotype were bulk fertilized, allowing for up to 110 potential crosses in each pure phenotype and 462 crosses in the site-wide control, excluding selfing. Larval survivorship was significantly higher at ambient temperatures (P_LRT_<0.01; Fig. S5). At the high temperature, all three phenotypes had significantly different survivorship trajectories (P_LRT_<0.01; Table S6, Fig. S5), with the highest survivorship in the site-wide cross phenotype (Fig. 3a). Larvae produced from bleached parents had substantially and significantly lower survivorship (4.9± 7.7%; mean ± 1SD) than larvae from nonbleached parents (24.9± 11.3%, ∼5-fold difference). Final survivorship averaged between 36.4-40.0% in the ambient treatments. Phenotype explained 96.3% of survivorship variance in high treatments (H^2^ = 0.963), and 17.3 % in ambient treatments (H^2^ = 0.173).

In 1.5 month-old juveniles, experiment-wide survivorship under ambient conditions was significantly higher than under heat stress (P_LRT_<0.01; Fig. S6), and the nonbleached phenotype in high temperatures had the highest overall survivorship. Nonbleached juveniles had higher survivorship than the site-wide cross (P_LRT_<0.01) or bleached phenotype (P_LRT_<0.01, ∼2-fold difference) under heat stress (Fig. 3b; Fig. S6). Further, nonbleached juvenile survivorship under temperature stress was not significantly different than either pure phenotype in ambient conditions (Fig. 3b; Table S7). Site-wide cross treatments had lower survivorship in both temperature treatments during the juvenile phase. Phenotype explained 92.9% of variance in high treatments (H^2^ = 0.929), and 0.2% in ambient treatments (H^2^ = 0.002).

*M. capitata* releases symbiont provisioned eggs which may contribute to downstream thermal tolerance, so we also quantified symbiont community over time in larvae using qPCR. Treatment did not have a significant main effect on symbiont community (beta-regression; *χ*^2^ =1.62, p=0.203). The symbiont communities were strongly differentiated between phentoypes (beta-regression; *χ*^2^ =146.6, p<0.001) and there was a significant interaction of phenotype, treatment and time (beta-regression; *χ*^2^ =13.2, p=0.040) with *Durusdinium* or *Cladocopium* dominating the nonbleached and bleached larvae, respectively (Fig. 3c). Nonbleached larvae typically had background levels of *Cladocopium*, but bleached larvae typically did not have any *Durusdinium*. The site-wide control cross has an approximately equal proportion of each genus at the initiation of heat stress, but a substantial decline in proportion *Durusdinium* occurred in the ambient treatments of the control cross (beta-regression, z-ratio=3.60, p=0.056); while this pattern was very similar in the high treatment, the only significant declines occurred between 0 and 44 hours (beta regression, z-ratio=6.04, p<0.001). The bleached (beta-regression, z-ratio <2.28, p>0.82) and nonbleached (beta-regression, z-ratio <1.41, p>0.99) crosses remain dominated by their initial symbiont communities.

The adaptive capacity of corals depends on a heritable, genomic response to thermal stress, so we used a select and re-sequence approach to define gene ontologies under heat selection that were not selected at ambient temperatures. We filtered the larval selection experiment to loci present in both the cross and nonbleached populations (n=31,794), which represents a realistic difference between ‘random’ breeding and selective breeding and calculated the Population Branch Statistic. This metric defines selection on a single population using two outgroups, in this case selection on final heat-stressed larvae compared to initial and the parental pool. In this way PBS minimizes differences between parents and recently fertilized larvae and minimizes sequencing artifacts. There was a significant relationship in selection strength (PBS) on individual loci between each pool (Fig. S7; lm p<0.001, R^2^ =0.335). Mean PBS was 1.5-fold higher in the cross treatment than the nonbleached treatment (Wilcox p<0.001).

Gene Ontology ranks (from GO_MWU) summarize the selection on a functional ontology under heat stress (using PBS), so we compared cross and nonbleached ranks to evaluate how selective breeding relates to random crosses representative of natural reproduction. Ranks of gene ontologies were significantly correlated in the global dataset (Fig. 4a gray; lm p<0.001, R^2^ =0.178) and were substantially more highly correlated among the loci which were significantly selected in both phenotypes (Fig. 4a orange; n=25, lm p<0.001, R^2^ =0.851). There were 119 and 90 ontologies significantly enriched for selected loci in the cross and nonbleached phenotypes, respectively, of which 23 were shared (Fig. 4b). Cell-signaling, immune response, DNA maintenance and repair, transcriptional regulation, cytoskeletal components and inflammation response were significantly enriched (FDR p<0.1) in both phenotypes for loci with high selection values under heat stress (Fig 4b). We also calculated enrichment in the difference between selection in nonbleached and cross larvae, respectively, to isolate the genomic consequences of selective breeding. Eighteen functional categories were significantly enriched, including antioxidant activity (GO:0016209;GO:0004601;GO:0016684) and protein Serine/Threonine/Tyrosine Kinase activity (GO:0004712) (Fig. 3a blue points, Table S8).

**Figure 4.**
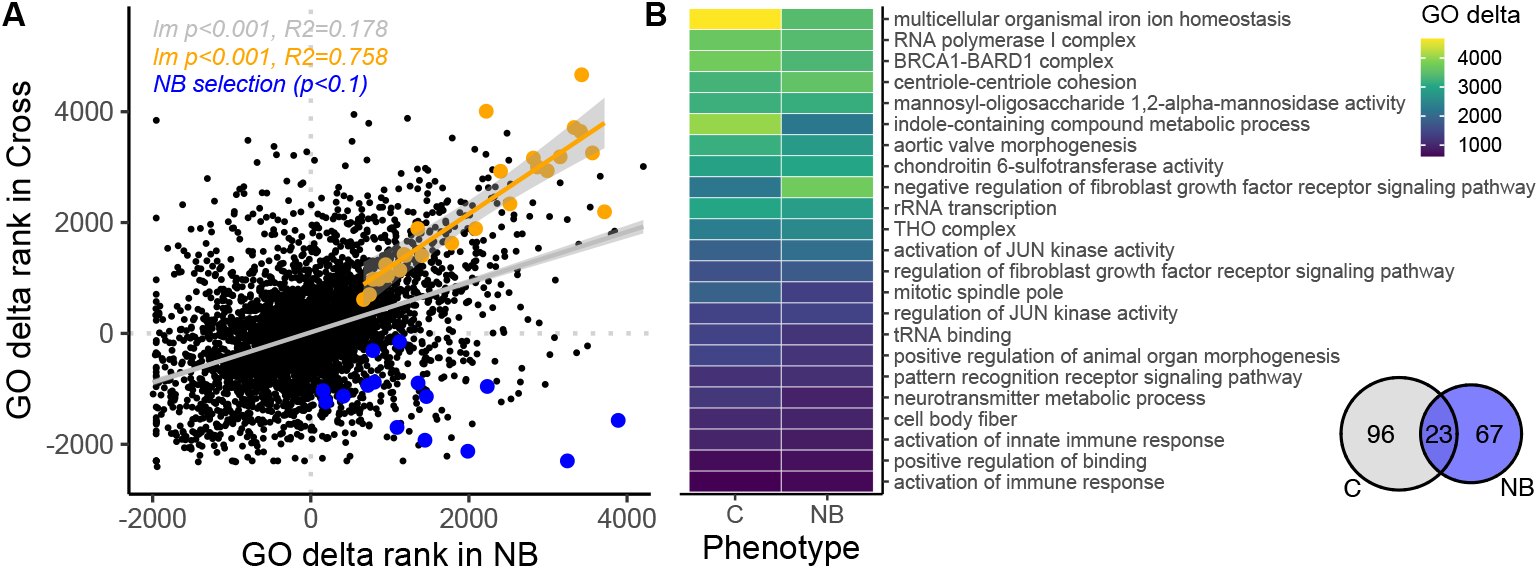
Selection Dynamics in Larval heat stress experiment. A) GO delta rank of cross and nonbleached larval pools after heat stress. High values represent gene ontologies with large PBS statistics. Orange points represent the 23 gene ontologies that were significant (FDR p<0.1) in both cross and nonbleached phenotypes, with lm best fit line. Black points represent the ontologies that were not significant in both treatments, with gray lm best fit line. Blue points represent gene ontologies with significant PBS differences between Nonbleached and Cross phenotypes, isolating the genomic effect of selective breeding. B) gene ontologies that were significantly enriched for large PBS values in both cross and nonbleached phenotypes. Venn diagram shows shared significant ontologies.

## Discussion

The long-term persistence of coral reefs requires adaptive differences based on heritable variation to keep pace with changing climate. Here, we demonstrate the genomic basis of intrapopulation variance in thermal tolerance in the absence of differential selection and show that it can be leveraged for selective breeding, increasing thermal tolerance in larvae and juveniles produced from known thermally tolerant parents.

We document strong intrapopulation genomic differentiation which explains a substantial proportion (24%) of bleaching tolerance in *Montipora capitata* from a single reef in Kāneʻohe
 Bay, Hawaiʻi. Due to, or despite, this variation, symbiont communities also exhibit the expected concentration of *Durusdinium* in historically thermally tolerant corals in this ecosystem^21,22^. Many of the canonical heat-stress response pathways distinguish phenotypes in the absence of historical differential selection, providing evidence for the genomic foundation of phenotypic variation over very small scales and highlighting the adaptive capacity of an individual coral reef without the migration or input of additional, pre-adapted genetic variance. These ontologies relating to growth/development (GO:0005138;GO:0070851), cell-signaling (GO:0008138), ribosomal constituents (GO:0003735), GTPase activity (GO:0051020) and protein localization, modification and breakdown (GO:0006518, GO:0043043, GO:0043603) have been documented extensively in previous work on both genomic and transcriptomic impacts of thermal tolerance^49,50^. We documented a single cellular compartment, the late endosome which is part of the response to a broad variety of environmental stressors in corals^50^. Interestingly, mitochondrial effects were not apparent in our dataset, although they have previously been implicated in larval survivorship and selection^14^.

The patterns resolved here also illustrate the biochemical consequences of gene ontologies which define bleaching phenotypes, including protein modification (i.e., protein phosphatase activity) and cell-signaling related to stress (i.e., ERK1/2 cascade), providing functional context for the role of host genotype. Interestingly, the molecular classes correlated with genomic drivers of phenotype are decoupled from the metabolites which directly differentiate phenotype in a global analysis. For example, Roach et al.^23^ showed that DGCC lipids and steroids were disproportionately important for distinguishing phenotype using machine learning and that fatty acids, monoacylglycerides, nucleotides, steroids and xanthins showed phenotype-specific responses to thermal stress^23^. None of these molecular classes are significantly enriched in the metabolite modules that correlate with gene ontologies defining phenotype in this study. We hypothesize that this difference reflects the host-specific regulation of physiology and biochemical patterns, rather than broader molecular patterns due to symbionts or the interaction of symbionts and the host. Although the correlations between GO loadings and metabolite module abundance were not significant after multiple comparisons correction, the congruent response of many molecules with several ontologies suggests that this is a biologically meaningful pattern, representing an important link that defines the downstream consequences of genomic variants besides eQTLs^51^.

There was also a link between two loci strongly predictive of adult phenotype and amino acid modifications and metabolism, primarily via Betaine Glycine. Betaine Glycine is extremely common across the domains of life, abundant in corals^52,53^, and found in *Montipora capitata* Kāneʻohe Bay^52,54^. Betaine Glycine is an amino-acid osmolyte that has well-documented protective effects on plants during abiotic stress and BG metabolism and biosynthesis pathways are found in corals^55^and commonly targeted in transgenic crop research^56^. This molecule is particularly important for the interaction of light and other abiotic stressors, including in corals^53^, which is closely aligned with the photosensitive nature of coral bleaching under high temperatures. Under these conditions, the repair of the D1 protein of photosystem II in symbionts is impaired, but the accumulation of BG prevents this obstruction and thereby limits oxidative stress. BG may also counteract cellular signals that initiate HSP70 expression^57^, representing a novel pathway to thermal tolerance in corals. Because symbiont communities maternally inherited in *M. capitata*^58^ and appear fixed through time^22^, we hypothesize that this variant and its downstream metabolomic differences impact the ability of the host to harbor *Cladocopium* or *Durusdinium* in this population, potentially through impacts on light tolerance. Importantly, patterns of BG could be influenced by selection not on variants in GDMT/SDMT itself, but some other regulatory machinery in close proximity, including PAX-C, or one of the methyltransferases, which could produce the same effects.

We also resolve the utility of ecologically realistic selective breeding in this system, which shows meaningful increases in survivorship after a single generation, especially once corals reach the juvenile stage. Breeding for heat tolerant larvae did not result in significant declines in survival at ambient temperatures, meaning there are minimal tradeoffs for these traits within our experiment, which is an important consideration in any breeding program. In addition, juvenile survivorship at high temperatures was not different from ambient survivorship in the nonbleached phenotype, suggesting that standing intrapopulation variance has the adaptive range to cover temperature increases like those tested here (+2.5°C C), rather than simply mute the declines expected under future conditions.

The positive effects of breeding reflect selection on diverse functional genes. Iron homeostasis and ferric iron binding were documented in the larval selection and adult association datasets, respectively, supporting a role for iron limitation in the governance of symbiosis and coral bleaching^59,60^. MAP kinase cascades were also important for adults and larval selection, though through complementary ontologies (ERK 1/2 cascades and JUN-kinase activity), highlighting the prominence of signaling pathways in coping with bleaching stress at various life history stages. In larvae, we also documented similar strong selection in both cross and nonbleached phenotypes on the BRCA1-BARD1 complex, which is involved in DNA repair and ubiquitination, rRNA transcription, and potential interactions between JUN-kinase activity and the THO complex, which regulates HSP70 export from the nucleus^61,62^. Importantly, the selection on the two crosses is broadly different, with less than a quarter of significantly selected ontologies shared between phenotypes, which indicates differential selection between the site-wide cross and nonbleached pool is much broader than coordinated selection. The weak but significant explanatory relationship between GO ranks in all ontologies (∼17%) suggests there is still a conserved response between phenotypes which allows for the decoupling of specific ontologies important for thermal tolerance.

We used the difference in PBS between nonbleached and cross treatments as a metric to isolate the genomic effect of selective breeding and examined functions that were enriched in high values. These functions included additional protein modification pathways (GO:0038083, GO:0004712), pH and ionic transport related ontologies (GO:0051453, GO:0051592, GO:2001225) and antioxidant/peroxidase activity (GO:0016209, GO:0004601), which is critical for dealing with stress related to reactive oxygen species during bleaching^63^. Overall, the heat selection of coral larvae shares some core functions across this population, while others are specific to the genetic input. This pattern highlights how selective breeding may influence thermal tolerance in *M. capitata* through the interaction of host genetics and symbiosis state.

We document broad-sense heritability >0.9 in both juveniles and larvae under thermal stress, but substantially lower estimates under ambient conditions, which supports the absence or weakness of tradeoffs in this system. The larval estimates are much higher than calculations from individual coral crosses from a single reef in the Persian Gulf ^64^ and estimates from more sparse parental input across latitudinal gradients ^14^. However, our broad-sense heritability calculations of juvenile survivorship closely mirror patterns at similar temperatures in *A. spathulata ^12^*, and we hypothesize that high heritability in this experiment corals is an additive effect of host and symbiont communities, where the more thermally tolerant *Durusdinium* dominant corals covary with targeted host-derived differences that cumulatively define most of the thermal capacity of the coral. Such high values can contribute to rapid adaptation on reefs^65^ and mean that thermally tolerant maternal transmitting corals may have an adaptive advantage under climate change.

The role of different symbionts in defining thermal tolerance has been documented in many studies in adults, but most corals have azooxanthellate larvae, so the role of infection (or transmission) of different symbionts in the larval stage remains poorly defined^66^, although these differences impact physiology^67^ and can dramatically alter host gene expression^68,69^. Juvenile *Acropora spathulata* infected with *Durusdinium trenchii* bleached less than those infected with other symbionts^70^, especially when one or both parents were from warm-adapted reefs, which closely mirrors survivorship patterns in juveniles in our study, assuming the *Cladocopium:Durusdinium* ratios found in larvae were maintained. While this system does not allow us to isolate host effects, the phenotypic variance explained by genotype and selection against *Durusdinium* in larvae strongly suggests that host and symbiont affects both impact thermal tolerance in this system.

Our results have profound implications for selective breeding and maintenance of adaptive potential, showing that the addition of tolerant corals within a population raises population-wide larval survivorship beyond the baseline of susceptible individuals. These results support the benefits of latitudinal crosses for increasing thermal tolerance^12,14,70^ and the importance of natural or assisted introduction of thermally tolerant stock for the persistence of reefs^10,71^. Tradeoffs in ambient conditions representative of the contemporary environment are minimal, with little difference in larval or juvenile survivorship between selectively bred tolerant corals and site-wide controls. Likewise, bleaching history had little explanatory power under ambient conditions (h<0.2) and we documented no differences in growth between these larval pools at ambient conditions. Selection of diverse, tolerant stocks for restoration, assisted migration or assisted gene flow is critical to the effectiveness of interventions^72^, and we show that this process can create meaningful changes over short time-scales, which may help offset crises of scale^10,11^.

The data presented here document the genomic basis of thermal tolerance and symbiosis state in corals upon which no differentiating selection has acted. This variance, which relates to intrapopulation differences in bleaching response, is the raw material upon which adaptation under climate change will act if previously bleached corals are the first to succumb to repeated bleaching events. Our data also suggest that the raw materials for selective breeding or natural adaptive recombination need not occur over large environmental gradients and that demographic processes generate this variance on individual reefs.

## Acknowledgments

We would like to thank Elizabeth Lenz for input on fertilization and rearing strategies in *Montipora capitata*, Katie Barott for initial colony tagging/surveys, Ingrid Knapp for library preparation advice, Justin Greer, Eva Majerová and Ariana Huffmyer for constructive comments on the manuscript and Kira Hughes, Chris Suchocki, Michelle Harangody and the Coral Resilience Lab for field and laboratory spawning assistance. Coral fragments and gamete bundles were collected under Hawaii DLNR permit SAP 2018-03 to HIMB. We gratefully acknowledge the technical support and advanced computing resources from University of Hawaii Information Technology Services – Cyberinfrastructure. This work was funded by a Paul G. Allen Family Foundation grant to Ruth D. Gates.

## Author Contributions

**CD** conceived and designed the study, collected and analyzed data and wrote the manuscript, **JH, TR** collected data, **NB, CH, CM, RAQ, JH** collected and analyzed data**, RDG** provided materials and resources. All authors edited and approved the final manuscript.

## Data Availability

All analysis scripts, ecological data and processed sequencing data will be made available at github.com/druryc/mcap_bleaching. Raw sequencing data is available at NCBI PRJNA597077.

