## Supplemental Figures and Tables for "Intrapopulation adaptive variance supports selective breeding in a reef-building coral"

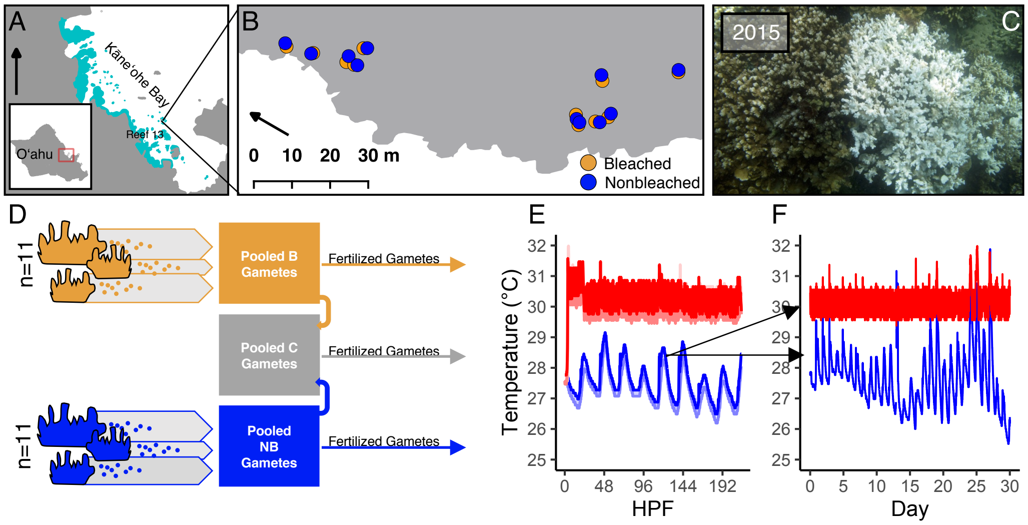


**Supplemental Figure 1 – Site and Experimental Framework**

A) Gamete collections were made at Reef 13 in Kāneʻohe Bay, Oʻahu, Hawaiʻi on 13 July 2018. B) Individual colonies were netted for collections using C) pairs identified in the 2015 bleaching event to minimize microhabitat differences and select for parental thermal tolerance. D) Gametes were collected from 11 colonies of each phenotype and used to create pools of all nonbleached gametes and all bleached gametes. The site-wide cross was then created from equal volumes of the bleached and nonbleached pools and all three pools were exposed to E) larval temperature treatments downstream starting 12 hours after fertilization.  At 109 hours after fertilization, larvae from all three phenotypes were allowed to settle on preconditioned aragonite plugs for 8 days and then exposed to F) juvenile temperature treatments. See Supplementary Fig. 1-2 for full details. Color scheme maintained in subsequent figures.

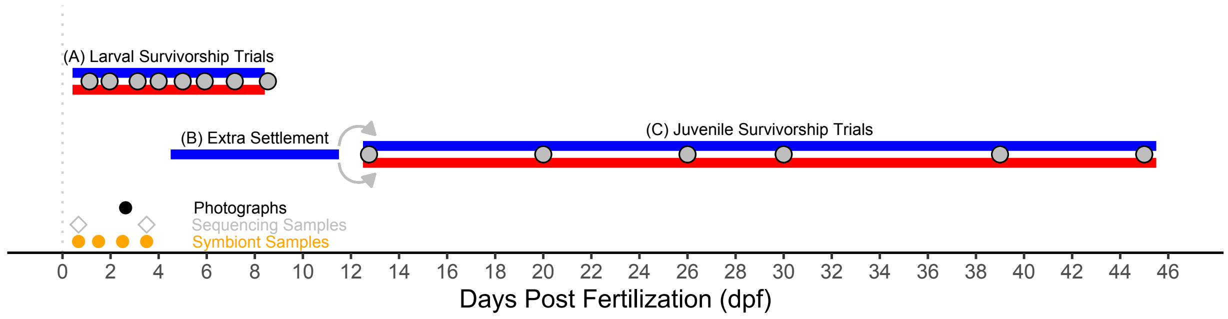


**Supplementary Figure 2 - Experimental Timeline**

Timeline in days post fertilization detailing temperature treatments, sequence, and data collection. Blue and red bars represent duration of treatment for larval and juvenile stress tests, with nested gray dots at survivorship survey timepoints. Points correspond to data collection for symbiont samples, larval sequencing and photographs for growth measurements. Briefly, (A) larvae were aliquoted into 50mL tubes for survivorship analysis at two temperatures on 1 dpf. Sequencing samples were collected from larval cultures (not from survivorship aliquots) at 1 and 4 dpf. (B) After final sequencing samples were collected (4 dpf) extra larvae at ambient temperatures were allowed to settle at ambient temperatures without interrupting survivorship aliquots. (C) These settled juveniles were randomly allocated to high and ambient temperatures at 12 dpf and monitored until 45dpf.

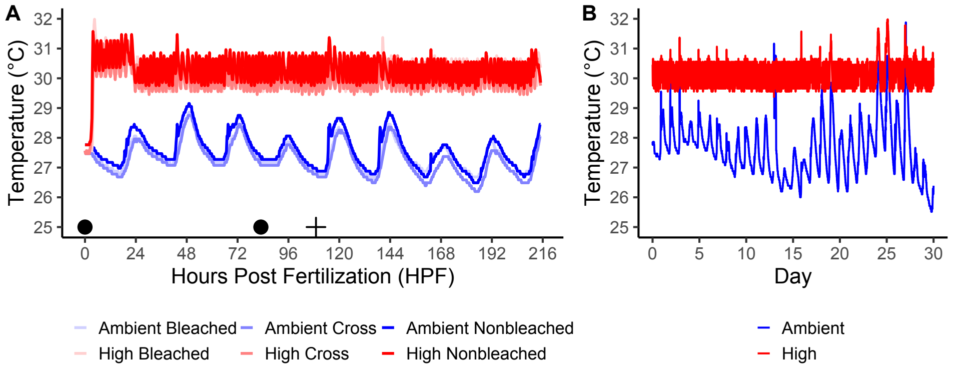


**Supplementary Figure 3 – Temperature Profiles**

A) Temperature profiles from each conical during the larval phase. Black circles denote timing of genetics sampling. Cross denotes timing of transfer of remnant ambient larvae to settlement chambers. B) Temperature profiles from each conical during the juvenile phase. Colors correspond to temperature treatment (blue=Ambient, red=High).

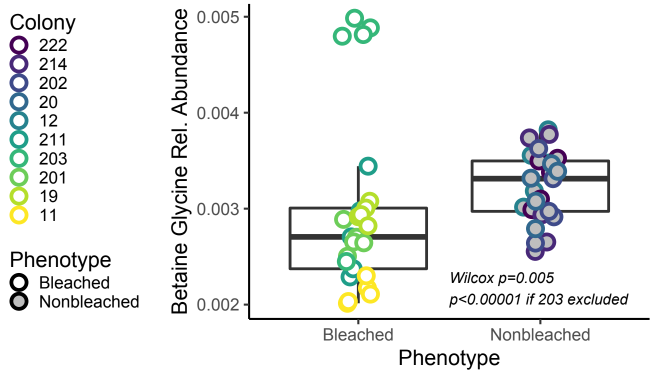


**Supplemental Figure 4 – Betaine Glycine Abundances**

Betaine Glycine abundances from parent colonies in this study.

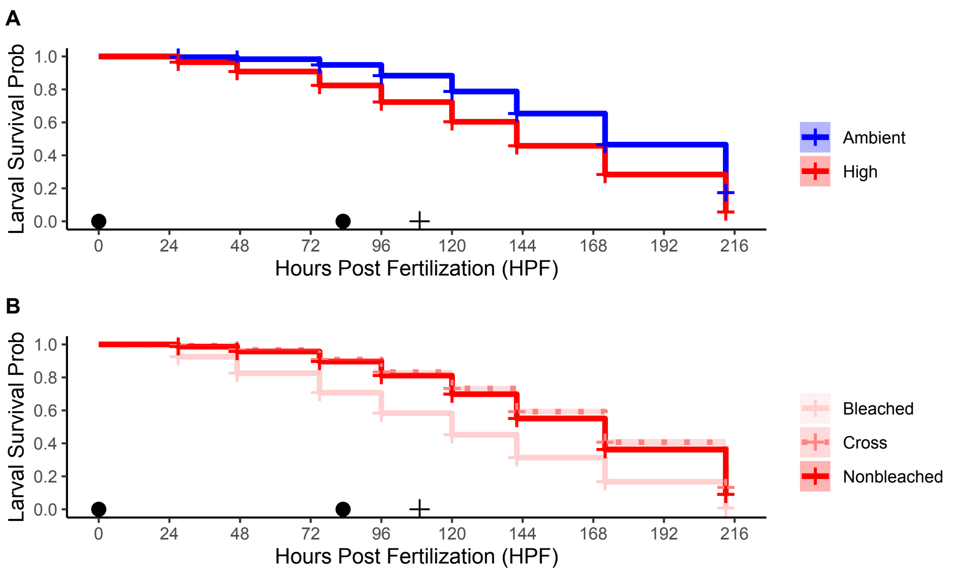


**Supplementary Figure 5 – Survivorship Dynamics of Larval *M. capitata***
A) Probability estimates from Kaplan-Meier larval survivorship fits for each temperature over hours in temperature treatment, line shown with 95% confidence interval shading. B) Probability estimates from Kaplan-Meier larval survivorship fits for each phenotype in high temperature treatment, line is shown with 95% confidence interval shading. Black circles denote timing of genetics sampling. Plus denotes timing of transfer of remnant ambient larvae to settlement chambers. Colors correspond to temperature treatment (blue=Ambient, red=High), transparency corresponds to phenotype.

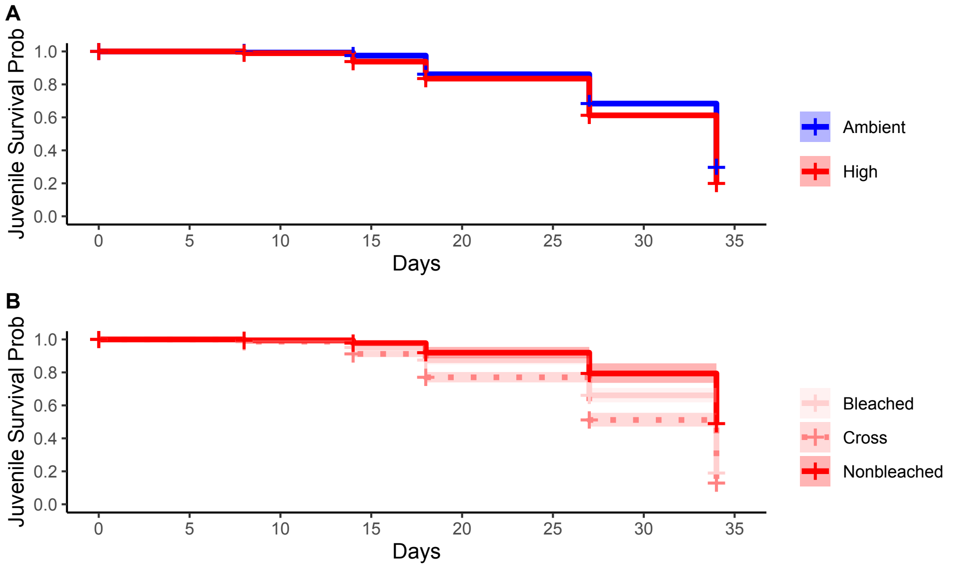


**Supplementary Figure 6 – Survivorship Dynamics of Juvenile *M. capitata***

A) Probability estimates from Kaplan-Meier juvenile survivorship fits for each temperature over hours in temperature treatment, line shown with 95% confidence interval shading. B) Probability estimates from Kaplan-Meier larval survivorship fits for each phenotype in high temperature treatment, line is shown with 95% confidence interval shading. Black circles denote timing of genetics sampling. Plus denotes timing of transfer of remnant ambient larvae to settlement chambers. Colors correspond to temperature treatment (blue=Ambient, red=High), transparency corresponds to phenotype.

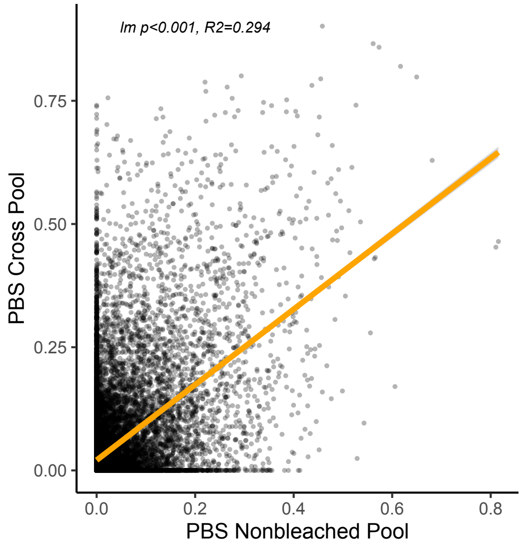


**Supplemental Figure 7**

Correlation of Population Branch Statistic (PBS) for the Cross and Nonbleached pool. PBS represents selection pressure under heat stress at each loci (n=31,794).

**Supplemental Table 1**

PBS calculation framework, comparing heat-selected larval pools of each phenotype to initial and parental pools. T= -log(1-$F_{ST}$), where pairwise $F_{ST}$ was calculated on a per-site basis for each variant (i.e., $T_{FA}$ = T based on pairwise $F_{ST}$

| Phenotype | (F) Focal Group | (A) Outgroup A | (B) Outgroup B | Formula |
|---|---|---|---|---|
| NB | Final Pool (n=250) | Initial Pool (n=250) | Nonbleached Parents (n=11) | PBS = $(T_{FA} + T_{FB} - T_{AB})/2$ |
| B | Final Pool (n=250) | Initial Pool (n=250) | Bleached Parents (n=10) | PBS = $(T_{FA} + T_{FB} - T_{AB})/2$ |
| Cross | Final Pool (n=250) | Initial Pool (n=250) | All Parents (n=21) | PBS = $(T_{FA} + T_{FB} - T_{AB})/2$ |

**Supplemental Table 2**

SIRIUS 4 parameters used for the *in silico* molecular formula annotations of the mass spectrometry data generated from MZmine 2. Mascot graphical format file (mgf) was imported and processed as follows:

| Module | Parameters | |
|---|---|---|
| Sirius | Instrument | Orbitrap |
| | $MS^2$ isotope scorer: | Score |
| | $MS^2$ mass deviation | 5 ppm |
| | Candidates | 5 |
| | Databases | Bio database, GNPS, Natural Products |
| ZODIAC | Default | |
| CSI: FingerID | Databases | Bio database, GNPS, Natural Products |
| | Adducts | $M+H]^+$, $[M+Na]^+$ and $[M+K]^+$ |
| | Enforce | True |
| CANOPUS | True | |

**Supplemental Table 3**

Annotations from individual genes significantly ($p<0.01$) associated with bleaching phenotype in adults. Black text denotes annotations from megablast, blue text denotes annotations from discontinuous megablast.

| locus | gene name | e-value | organism | function | ref |
|---|---|---|---|---|---|
| 114:270077-1166526(+)_382645 | Ethanolamine phosphotransferase 1-like | 2e-64 | *Acropora digitifera* | Sphingolipid biosynthesis | 25 |
| 1142:22067-267875(+)_240011 | Ankyrin repeat and zinc-finger domain-containing protein 1-like | 2e-12 | *Acropora digitifera* | Immunity | 1,32 |
| 135:165149-681565(+)_107553 | Monocarboxylate transporter 10-like | 1e-124 | *Acropora millepora* | Nitrogen cycling; amino acid metabolism | 2 |
| 16:103141-1930322(+)_88237 | Histamine H2 receptor-like | 3e-71 | *Acropora millepora* | Immune response | 3 |
| 175:84843-855127(+)_390354 | PAX-C | 4e-114 | *Montipora sp.* | Growth/development; host-algal symbiosis regulation | 4,5 |
| 23:0-1877333(+)_1275116 | Neuropeptide SIFamide receptor | 8e-37 | *Orbicella faveolata* | Larval migration, settlement, metamorphosis | 6,7 |
| 2374_33003 | 18s rRNA | 0.0 | *Montipora verrucosa* | Protein synthesis | 28 |
| 2644:0-29873(+)_8082 | Histamine N-methyltransferase-like | 0.0 | *Acropora millepora* | Methylation (humans) | 8 |
| 316:172135-793760(+)_405758 | PAX-C | 6e-83 | *Montipora solanderi* | Growth/development, host-algal symbiosis regulation | 4,5 |
| 32:221614-1703322(+)_986481 | Histamine H2 receptor-like | 2e-38 | *Acropora millepora* | Immune response | 3 |
| 362:154151-476426(+)_296306 | Unknown | 2e-24 | *Acropora millepora* | Unknown | - |
| 416:25677-496747(+)_417623 | Mucin-like | 2e-31 | *Acropora digitifera* | Mucus generation/secretion; DNA damage repair | 9,23 |
| 462:90630-579403(+)_429238 | Octopamine receptor beta-1R-like | 3e-73 | *Orbicella faveolata* | Oocyte development/maturation | 10 |

| | | | | | |
|---|---|---|---|---|---|
| 5:218022-2430108(+)_851904 | Adenosine receptor A2a-like | 2e-93 | *Acropora millepora* | Immune response; inflammation | 1,11 |
| 50:672149-1522693(+)_465625 | Polyubiquitin-C | 0.0 | *Stylophora pistillata* | Oxidative stress resistance; protein catabolism | 12,29 |
| 553:474523-500157(+)_13812 | Ribonuclease Y-like | 7e-133 | *Orbicella faveolata* | Unfolded protein response | 13 |
| 615:456522-469093(+)_1880 | Plexin-B-like | 1e-47 | *Acropora millepora* | Cytoskeleton dynamics; cell adhesion; axon guidance | 14,23 |
| 69:44661-1355009(+)_72740 | Phosphoinositide 3-kinase regulatory subunit 4-like | 2e-58 | *Acropora digitifera* | Host autophagy | 15 |
| 94:79413-1187499(+)_1064347 | kelch-like protein diablo | 8e-113 | *Acropora millepora* | Oxidative stress response; protein ubiquination | 26, 30 |
| chr1_RagTag_22018874 | Golgi-associated PDZ and coiled-coil motif-containing protein-like | 8e-77 | *Acropora millepora* | Protein binding/transport | 16 |
| chr1_RagTag_31003888 | Dynein assembly factor 3, axonemal-like | 2e-32 | *Actinia tenebrosa* | Cytoskeletal construction; autophagy | 17,31 |
| chr10_RagTag_8284035 | Putative ankyrin repeat protein RF_0381 | 1e-49 | *Dendronephthya gigantea* | Immunity | 1 |
| chr11_RagTag_4203672 | Neurexin | 2e-68 | *Acropora digitifera* | Biomineralization | 18 |
| chr11_RagTag_4215008 | TFIID subunit 5-like | 1e-46 | *Acropora digitifera* | Cell signaling; transcription factor activity; protein binding | 21 |
| chr12_RagTag_25257908 | Phosphoinositide 3-kinase regulatory subunit 4-like | 5e-65 | *Acropora digitifera* | Host autophagy | 15 |
| chr2_RagTag_20154540 | Tetratricopeptide repeat protein 28 | 2e-108 | *Orbicella faveolata* | Apoptotic signaling | 27 |
| chr4_RagTag_11047240 | kelch-like protein diablo | 6e-45 | *Acropora digitifera* | Oxidative stress response; protein ubiquination | 26,30 |

| | | | | | |
|---|---|---|---|---|---|
| chr4_RagTag_5271861 | Unknown | 0.0 | *Acropora millepora* | Unknown | - |
| chr5_RagTag_11207344 | Unknown | 0.0 | *Acropora millepora* | Unknown | - |
| chr6_RagTag_13411819 | Arginine/serine-rich protein PNISR-like | 8e-145 | *Acropora digitifera* | Protein synthesis | 19 |
| chr8_RagTag_12409112 | Neurexin | 6e-70 | *Acropora digitifera* | Biomineralization | 18 |
| chr8_RagTag_4922595 | Ankyrin repeat and zinc-finger domain-containing protein 1-like | 2e-12 | *Acropora millepora* | Immunity | 1,32 |
| Sc0000015_RagTag_1507093 | E3 ubiquitin-protein ligase MARCH5-like | 2e-58 | *Acropora millepora* | *Symbiodiniaceae* stress response; protein degradation | 20,21 |
| Sc0000134_RagTag_502413 | Somatostatin receptor type 4-like | 3e-160 | *Acropora millepora* | Myoregulatory activity; neural pathways; MAP kinase and adenylate cyclase interactions | 22 |
| Sc0000201_RagTag_77657 | D-inositol 3-phosphate glycosyltransferase-like | 4e-50 | *Acropora digitifera* | Host/algal symbiosis regulation | 23 |
| Sc0000204_RagTag_980637 | Plexin-B-like | 1e-64 | *Acropora millepora* | Cytoskeleton dynamics; cell adhesion; axon guidance | 14,23 |
| xpSc0000570_RagTag_25400 | Cationic amino acid transporter 1-like | 2e-58 | *Stylophora pistillata* | Protein/membrane formation | 24 |

**Supplemental Table 4**

Summary of Gene Ontologies enriched in significantly high LRT values, suggesting functions that are significantly different between adult phenotypes and distinguish bleaching tolerance.

| Term | Name | P (fdr) | Category |
|---|---|---|---|
| GO:0006412 | translation | 0.084 | BP |
| GO:0006476; GO:0035601; GO:0098732 | protein deacetylation | 0.028 | BP |
| GO:0006518 | peptide metabolic process | 0.046 | BP |
| GO:0006575 | cellular modified amino acid metabolic process | 0.045 | BP |
| GO:0018200 | peptidyl-glutamic acid modification | 0.028 | BP |
| GO:0034334; GO:0034332 | adherens junction organization | 0.045 | BP |
| GO:0043043 | peptide biosynthetic process | 0.059 | BP |
| GO:0043603 | cellular amide metabolic process | 0.085 | BP |
| GO:0070373 | negative regulation of ERK1 and ERK2 cascade | 0.048 | BP |
| GO:0071333; GO:0009749; GO:0071331; GO:0009746; GO:0034284; GO:0009743; GO:0071326; GO:0071322 | cellular response to hexose stimulus | 0.077 | BP |
| GO:0090630 | activation of GTPase activity | 0.085 | BP |
| GO:1901565 | organonitrogen compound catabolic process | 0.077 | BP |
| GO:1902414 | protein localization to cell junction | 0.024 | BP |
| GO:1903614; GO:1903613; GO:1904894; GO:0018279; GO:0006487; GO:0018196 | protein N-linked glycosylation | 0.027 | BP |
| GO:0031902 | late endosome membrane | 0.099 | CC |
| GO:0000287 | magnesium ion binding | 0.094 | MF |
| GO:0003735 | structural constituent of ribosome | 0.094 | MF |
| GO:0003887; GO:0004523; GO:0016891 | DNA-directed DNA polymerase activity | 0.094 | MF |
| GO:0004177; GO:0008238; GO:0070006; GO:0008235 | exopeptidase activity | 0.094 | MF |

| | | | |
|---|---|---|---|
| GO:0004190; GO:0070001 | aspartic-type endopeptidase activity | 0.094 | MF |
| GO:0008138 | protein tyrosine/serine/threonine phosphatase activity | 0.093 | MF |
| GO:0008199; GO:0004322; GO:0016724; GO:0016722 | ferric iron binding | 0.094 | MF |
| GO:0016758; GO:0016757 | transferase activity, transferring glycosyl groups | 0.093 | MF |
| GO:0017016; GO:0031267 | small GTPase binding | 0.094 | MF |
| GO:0017048 | Rho GTPase binding | 0.094 | MF |
| GO:0030144; GO:0140103; GO:0030145; GO:0004864; GO:0019212 | protein phosphatase inhibitor activity | 0.093 | MF |
| GO:0051020 | GTPase binding | 0.028 | MF |
| GO:0070851; GO:0005138 | growth factor receptor binding | 0.094 | MF |

**Supplemental Table 5**
Summary of Gene Ontologies enriched in significantly low LRT values, suggesting functions that are significantly similar between adult phenotypes and do not distinguish bleaching tolerance.

| Term | Name | P (fdr) | Category |
|---|---|---|---|
| GO:0001667 | ameboidal-type cell migration | 0.048 | BP |
| GO:0001932; GO:0042325 | regulation of phosphorylation | 0.037 | BP |
| GO:0002252; GO:0051607; GO:0009615 | immune effector process | 0.045 | BP |
| GO:0002376 | immune system process | 0.077 | BP |
| GO:0008037 | cell recognition | 0.077 | BP |
| GO:0008202 | steroid metabolic process | 0.006 | BP |
| GO:0009605 | response to external stimulus | 0.077 | BP |
| GO:0009888 | tissue development | 0.070 | BP |
| GO:0009952 | anterior/posterior pattern specification | 0.051 | BP |
| GO:0010035 | response to inorganic substance | 0.045 | BP |
| GO:0010038 | response to metal ion | 0.084 | BP |
| GO:0010557; GO:0010628; GO:0031328; GO:0009891 | positive regulation of gene expression | 0.054 | BP |
| GO:0010638 | positive regulation of organelle organization | 0.048 | BP |
| GO:0016192 | vesicle-mediated transport | 0.027 | BP |
| GO:0019827; GO:0098727 | stem cell population maintenance | 0.048 | BP |
| GO:0022604 | regulation of cell morphogenesis | 0.056 | BP |
| GO:0030278 | regulation of ossification | 0.077 | BP |
| GO:0030517 | negative regulation of axon extension | 0.095 | BP |
| GO:0031122 | cytoplasmic microtubule organization | 0.070 | BP |
| GO:0043207; GO:0009607; GO:0051707 | response to biotic stimulus | 0.029 | BP |
| GO:0045087 | innate immune response | 0.045 | BP |
| GO:0045596 | negative regulation of cell differentiation | 0.044 | BP |
| GO:0045597 | positive regulation of cell differentiation | 0.092 | BP |
| GO:0045859; GO:0043549; GO:0051338 | regulation of transferase activity | 0.008 | BP |
| GO:0045926 | negative regulation of growth | 0.083 | BP |
| GO:0045944 | positive regulation of transcription by RNA polymerase II | 0.045 | BP |
| GO:0050685 | positive regulation of mRNA processing | 0.085 | BP |

| | | | |
|---|---|---|---|
| GO:0050793 | regulation of developmental process | 0.056 | BP |
| GO:0051093 | negative regulation of developmental process | 0.024 | BP |
| GO:0051094 | positive regulation of developmental process | 0.070 | BP |
| GO:0051128 | regulation of cellular component organization | 0.020 | BP |
| GO:0051130 | positive regulation of cellular component organization | 0.028 | BP |
| GO:0051240 | positive regulation of multicellular organismal process | 0.026 | BP |
| GO:0051241 | negative regulation of multicellular organismal process | 0.027 | BP |
| GO:0051704 | multi-organism process | 0.045 | BP |
| GO:0051965; GO:0051963 | regulation of synapse assembly | 0.085 | BP |
| GO:0070988 | demethylation | 0.029 | BP |
| GO:0070989 | oxidative demethylation | 0.006 | BP |
| GO:0071360; GO:0071359; GO:0009597; GO:1900246; GO:0039535; GO:0039531; GO:0032481 | cellular response to dsRNA | 0.024 | BP |
| GO:0090068; GO:0045787 | positive regulation of cell cycle | 0.084 | BP |
| GO:0005783 | endoplasmic reticulum | 0.099 | CC |
| GO:0048786 | presynaptic active zone | 0.099 | CC |
| GO:1990909; GO:1990851 | Wnt signalosome | 0.099 | CC |
| GO:0005509 | calcium ion binding | 0.094 | MF |
| GO:0008395; GO:0070330; GO:0016712; GO:0032451; GO:0101020; GO:0008401; GO:0050649 | aromatase activity | 0.009 | MF |
| GO:0015026; GO:1904928; GO:0071936; GO:0042813 | coreceptor activity | 0.094 | MF |
| GO:0020037; GO:0046906 | tetrapyrrole binding | 0.094 | MF |
| GO:1990837; GO:0003690; GO:0043565 | double-stranded DNA binding | 0.093 | MF |

**Supplemental Table 6**

Results from pairwise comparisons of Likelihood Ratio tests between larval Kaplan-Meier survivorship fits for each Phenotype~Temperature combination. A=Ambient, H=High, B= Bleached, C= Site-Wide Cross, NB= Nonbleached. p<0.05*; p<0.01**; p<0.001***

| | AB | AC | ANB | HB | HC | HNB |
|---|---|---|---|---|---|---|
| Ambient Bleached | | | | | | |
| Ambient Cross | *** | | | | | |
| Ambient Nonbleached | ns | * | | | | |
| High Bleached | *** | *** | *** | | | |
| High Cross | *** | ** | *** | *** | | |
| High Nonbleached | *** | *** | *** | *** | *** | |

**Supplemental Table 7**

Results from pairwise comparisons of Likelihood Ratio tests between juvenile Kaplan-Meier survivorship fits for each Phenotype~Temperature combination. A=Ambient, H=High, B= Bleached, C= Site-Wide Cross, NB= Nonbleached. $p<0.05$*; $p<0.01$**; $p<0.001$***

| | AB | AC | ANB | HB | HC | HNB |
|---|---|---|---|---|---|---|
| Ambient Bleached | | | | | | |
| Ambient Cross | *** | | | | | |
| Ambient Nonbleached | ns | *** | | | | |
| High Bleached | *** | ns | ** | | | |
| High Cross | *** | ** | *** | *** | | |
| High Nonbleached | ns | *** | ns | *** | *** | |

**Supplemental Table 8**

Summary of Gene Ontologies enriched in significantly high differences between PBS in nonbleached and cross corals, isolating functions that are selected significantly more strongly in nonbleached corals and highlighting effects of selective breeding.

| **Term** | **Name** | **P (fdr)** | **Category** |
|---|---|---|---|
| GO:0032230 | positive regulation of synaptic transmission, GABAergic | 0.035 | BP |
| GO:0032849; GO:0032847 | regulation of cellular pH reduction | 0.035 | BP |
| GO:0004712 | protein serine/threonine/tyrosine kinase activity | 0.040 | MF |
| GO:0043812; GO:0034596 | phosphatidylinositol phosphate 4-phosphatase activity | 0.040 | MF |
| GO:0015491; GO:0015298 | cation:cation antiporter activity | 0.054 | MF |
| GO:0008307 | structural constituent of muscle | 0.062 | MF |
| GO:0016209; GO:0004601; GO:0016684 | antioxidant activity | 0.062 | MF |
| GO:0022821; GO:0008273 | potassium ion antiporter activity | 0.083 | MF |
| GO:0032160 | septin filament array | 0.084 | CC |
| GO:0097733 | photoreceptor cell cilium | 0.084 | CC |
| GO:0150051 | postsynaptic Golgi apparatus | 0.084 | CC |
| GO:0006900 | vesicle budding from membrane | 0.090 | BP |
| GO:0032228 | regulation of synaptic transmission, GABAergic | 0.090 | BP |
| GO:0038083 | peptidyl-tyrosine autophosphorylation | 0.090 | BP |
| GO:0051453; GO:0030641; GO:0006885; GO:0030004 | regulation of pH | 0.090 | BP |
| GO:0051592 | response to calcium ion | 0.090 | BP |
| GO:0060402; GO:0060401; GO:0097553 | cytosolic calcium ion transport | 0.090 | BP |
| GO:2001225 | regulation of chloride transport | 0.090 | BP |
